# Pemphigus vulgaris autoantibodies induce an ER stress response

**DOI:** 10.1101/2024.08.22.608849

**Authors:** Coryn L. Hoffman, Navaneetha Krishnan Bharathan, Yoshitaka Shibata, William Giang, Johann E. Gudjonsson, John T. Seykora, Stephen M. Prouty, Aimee S. Payne, Andrew P. Kowalczyk

**Author notes:** Correspondence: Andrew P. Kowalczyk Department of Dermatology Pennsylvania State University College of Medicine Hershey, PA 17033, USA. These authors contributed equally to this work (Coryn L. Hoffman, Navaneetha Krishnan Bharathan).

## Abstract

Desmosomes are intercellular junctions that mediate cell-cell adhesion and are essential for maintaining tissue integrity. Pemphigus vulgaris (PV) is an autoimmune epidermal blistering disease caused by autoantibodies (IgG) targeting desmoglein 3 (Dsg3), a desmosomal cadherin. PV autoantibodies cause desmosome disassembly and loss of cell-cell adhesion, but the molecular signaling pathways that regulate these processes are not fully understood. Using high-resolution time-lapse imaging of live keratinocytes, we found that ER tubules make frequent and persistent contacts with internalizing Dsg3 puncta in keratinocytes treated with PV patient IgG. Biochemical experiments demonstrated that PV IgG activated ER stress signaling pathways, including both IRE1⍺ and PERK pathways, in cultured keratinocytes. Further, ER stress transcripts were upregulated in PV patient skin. Pharmacological inhibition of ER stress protected against PV IgG-induced desmosome disruption and loss of keratinocyte cell-cell adhesion, suggesting that ER stress may be an important pathomechanism and therapeutically targetable pathway for PV treatment. These data support a model in which desmosome adhesion is integrated with ER function to serve as a cell adhesion stress sensor that is activated in blistering skin disease.

## INTRODUCTION

Desmosomes are intercellular junctions that link the intermediate filament cytoskeleton to sites of cell-cell contact (Bharathan et al., 2024 in press; Perl et al., 2024; Zimmer and Kowalczyk, 2024 in press). These junctions mediate robust cell-cell adhesion and are critical for integrity of tissues that experience high levels of mechanical stress, including the skin and heart (Müller et al., 2021; Toivola et al., 2024; Waschke, 2008). The desmosomal cadherins, desmogleins and desmocollins, form adhesive interactions that are coupled to the intermediate filament cytoskeleton by intracellular plaque proteins. These desmosomal plaque proteins include armadillo proteins such as plakoglobin and plakophilins, as well as plakin family proteins, including desmoplakin (Delva et al., 2009). Together, these proteins mechanically integrate cells within the epidermis (Perl et al., 2024). Desmosome disruption due to genetic defects, autoimmunity, or infection causes tissue fragility and is the hallmark of various skin diseases (Hegazy et al., 2022).

Pemphigus vulgaris (PV) is an autoimmune bullous disease in which autoantibodies (IgG) against desmoglein 3 (Dsg3) cause steric hindrance, Dsg3 internalization, desmosome disassembly, and loss of keratinocyte adhesion in mucosal epithelia and epidermal keratinocytes (Amagai and Stanley, 2012; Egu et al., 2022; Spindler et al., 2023; Stahley and Kowalczyk, 2015). Clinically, patients present with blisters on the skin and erosions on mucous membranes (Hammers and Stanley, 2020; Kridin and Schmidt, 2021). Current treatments for PV involve immunosuppression (Abeles et al., 2024; Ellebrecht et al., 2022; Ujiie et al., 2021), which puts patients at serious risk for infection. These concerns underscore the need for a deeper understanding of PV pathomechanisms that could lead to the development of more targeted therapies. While it is well-established that PV autoantibodies against Dsg3 cause desmosome disassembly and loss of keratinocyte adhesion, the signaling mechanisms by which PV IgG cause these pathogenic responses are not fully understood.

We recently reported the structure and dynamics of a novel endoplasmic reticulum (ER)- desmosome complex (Bharathan et al., 2023). The ER is involved in a multitude of cellular processes including protein biosynthesis, lipid synthesis and transfer among organelles, and maintenance of calcium homeostasis (Prinz et al., 2020; Schwarz and Blower, 2016; Voeltz et al., 2024). In addition, the ER is an important stress-sensing organelle that responds to various forms of cellular stress, including metabolic, chemical, and mechanical stresses (Hetz et al., 2020; Mauro, 2014; Phuyal et al., 2023). ER stress has been implicated in the pathogenesis of various skin diseases such as melanoma, Darier’s disease, rosacea, vitiligo, and epidermolysis bullosa simplex (Evtushenko et al., 2021; Park et al., 2019). Interestingly, PV IgG activate p38 MAPK, a stress-activated protein kinase that is known to be involved in ER stress signaling (Cipolla et al., 2017). However, a functional relationship between ER stress and desmosome integrity has yet to be elucidated in the context of PV.

Using live-cell time-lapse confocal microscopy, we find that ER tubules localize to desmosomes and remain associated with Dsg3 puncta during PV IgG-induced internalization. ER stress transcripts are upregulated in PV patient skin, and PV IgG rapidly activates ER stress signaling by both the IRE1⍺ and PERK pathways in cultured keratinocytes treated with PV IgG. Pharmacological inhibition of ER stress prevents PV IgG-induced desmosome disruption and loss of keratinocyte adhesion. These findings indicate that desmosomal adhesion is integrated with ER function to serve as a sensor of desmosomal adhesion stress and that ER signaling pathways could be a potential therapeutic target for the treatment of PV and related disorders.

## RESULTS

### ER tubules are at desmosomes during disassembly in PV

We recently reported an association of peripheral ER tubules with keratin filaments and the desmosome plaque (Bharathan et al., 2023). In the present study, we used time-lapse spinning disk confocal imaging to determine how the ER-desmosome complex dynamically responds during PV IgG-induced desmosome disassembly. We used lentivirus to generate N/TERT human keratinocyte cell lines (Dickson et al., 2000) stably expressing KDEL-StayGold to label the ER lumen (Hirano et al., 2022). Cell surface Dsg3 was labelled with a fluorescently conjugated non-pathogenic mAb (P2C2-CF568) (Cho et al., 2019) directed against the Dsg3 extracellular domain. In cells treated with normal human (NH) IgG, ER tubules stably associated with junctional Dsg3 and remained anchored several hours after IgG addition (Figure 1a, Supplementary Video 1). PV IgG-treated cells exhibited similarly stable ER-Dsg3 associations initially (Figure 1b, Supplementary Video 2). This initial stage was followed by stereotypical Dsg3 clustering at cell-cell contacts, and then formation of linear arrays perpendicular to the cell-cell contact, as described previously (Jennings et al., 2011). Remarkably, during these dynamic reorganization events, ER tubules maintained associations with both Dsg3 clusters and linear arrays (Figure 1c, Supplementary Video 3). These data demonstrate a stable structural association between ER tubules and desmosomes during PV IgG-mediated desmosome disassembly.

**Figure 1.**
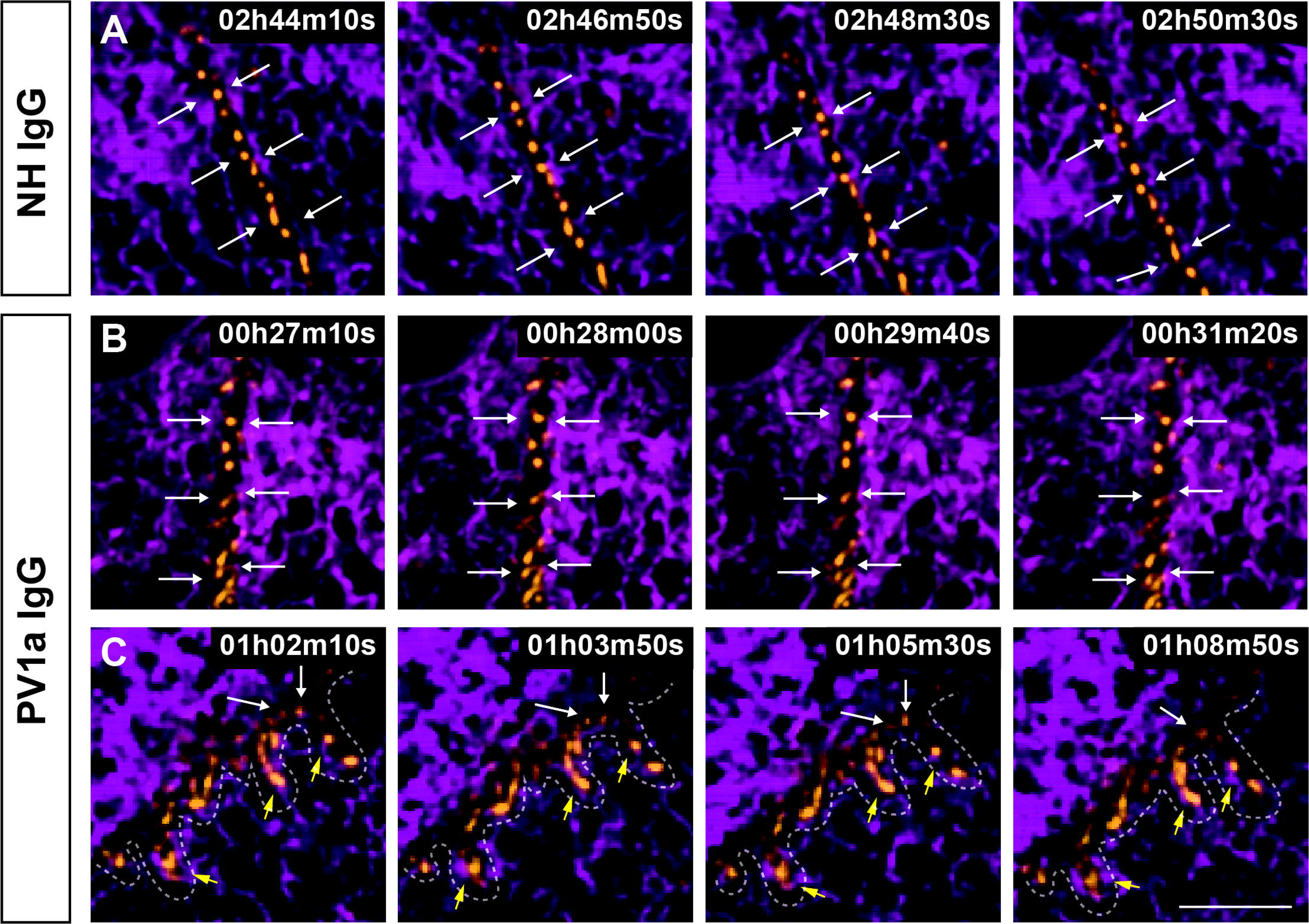
ER tubules localize to desmosomes and remain associated with Dsg3 puncta during PV IgG-induced internalization. N/TERT keratinocytes expressing luminal ER marker KDEL- StayGold (magenta) were pre-labeled with a fluorescently conjugated (CF568) Dsg3 antibody (orange), treated with IgG, and imaged every 10 sec for various lengths of time during the first 3.5 hours following IgG addition. **(a)** Dsg3 puncta formed stable associations with peripheral ER tubules in NH IgG-treated cells even 3 hours after IgG addition (white arrows). **(b)** Stable Dsg3- ER associations were seen within the first 30 minutes following addition of PV1a IgG (white arrows). **(c)** ER tubules formed stable associations with Dsg3 puncta undergoing clustering (white arrows) and wrapped around linear arrays (yellow arrows) forming perpendicular to the cell-cell contact (dashed line). Scale bar = 5 μm.

### PV IgG induce a rapid ER stress response

The physical association of ER with desmosomes during PV IgG-induced desmosome disassembly raised the possibility that ER signaling pathways are activated upon keratinocyte exposure to PV IgG. To determine if PV IgG induce ER stress signaling upon binding to Dsg3, we treated primary normal human epidermal keratinocytes (NHEKs) with PV IgG from several patients (PV1, PV2, and PV3) for 15 minutes to 1 hour and probed for IREIα and eIF2α phosphorylation as markers of ER stress pathway activation. Western blot analysis revealed that IRE1⍺ and eIF2⍺ phosphorylation were upregulated within 15 minutes of keratinocyte exposure to PV IgG (Figure 2a). To investigate if an ER stress response is observed in PV patients, we performed qRT-PCR of PV patient epidermis to evaluate the transcriptional levels of key ER stress markers. The expression of *DDIT3, XBP1s, ATF3,* and *HSPA5* were increased in PV patient skin biopsies from blister margins relative to normal healthy (NH) skin (Figure 2b).

**Figure 2.**
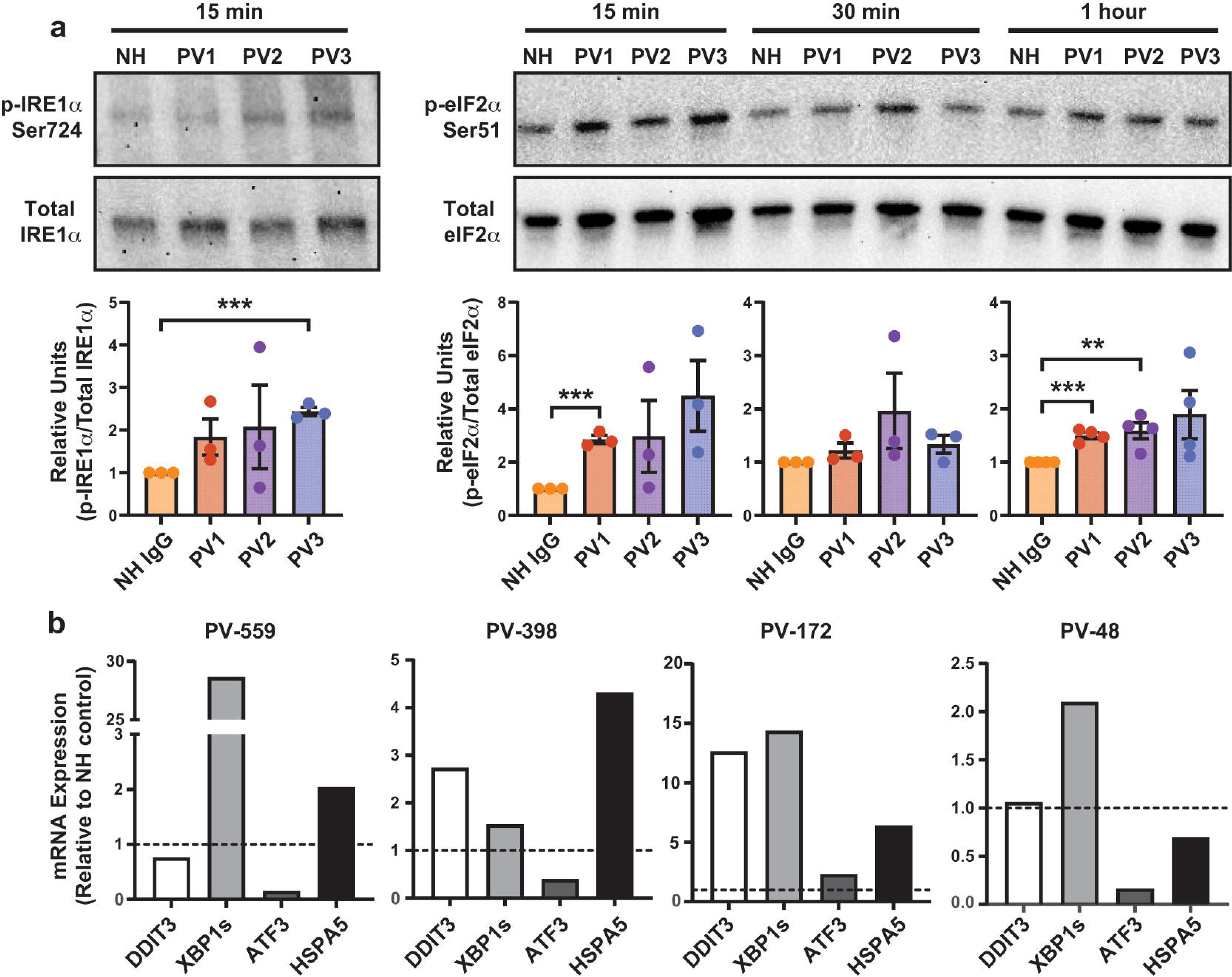
ER stress is activated in PV IgG-treated keratinocytes and PV patient skin. (a) Western blots showing ER stress marker phosphorylation in NHEKs treated with NH or PV IgG (PV1, PV2, or PV3) for 15 minutes to 1 hour. PV3 alone significantly increased IRE1⍺ phosphorylation after 15 min. PV1 alone significantly increased eIF2⍺ phosphorylation after 15 minutes and 1 hour. **(b)** Expression of ER stress transcripts was monitored by qRT-PCR in PV patient skin biopsies from blister margins. All four patients show differential upregulation of ER stress markers relative to NH controls (dashed line). Mean value ± SEM is represented. ** *P* < 0.01, and *** *P* < 0.001 using an unpaired student’s t-test comparing PV IgG-treated to NH IgG- treated.

While the relative expression of these transcripts was variable, at least one of these markers was upregulated in each of four different patient samples. Collectively, these data demonstrate that ER stress pathways are activated in both PV patient epidermis and *in vitro* cell culture models of PV.

### ER stress inhibitors prevent PV IgG-induced desmosome disruption and loss of adhesion in cultured keratinocytes

Previous studies indicate that desmosome disassembly and disruption of keratinocyte adhesion occur several hours after PV IgG binding (Calkins et al., 2006; Jennings et al., 2011). Since the ER stress response appears to be temporally upstream of desmosome disruption and loss of adhesion in PV, we investigated whether ER stress inhibition is protective against PV IgG-induced desmosome disruption. We inhibited ER stress with miglustat, a pharmacological chaperone (Abian et al., 2011; Alfonso et al., 2005; Savignac et al., 2014), or GSK2606414, a PERK inhibitor (Axten et al., 2012). To determine if miglustat and GSK2606414 prevent the PV IgG-induced ER stress response, we treated NHEKs with either miglustat or GSK2606414, followed by exposure to either NH or PV IgG for 15 minutes to 1 hour and probed for phosphorylated ER stress markers. Western blot analysis revealed that treatment of keratinocytes with PV1, PV2, or PV3 significantly increased IRE1⍺ and eIF2⍺ phosphorylation within 15 minutes. Miglustat pre-treatment abrogated IRE1⍺ phosphorylation by PV autoantibodies (Figure 3a) but did not reduce eIF2⍺ phosphorylation (Figure 3b). Further, GSK2606414 significantly reduced IRE1⍺ phosphorylation induced by the PV3 antibody, but not by PV1a or PV2 (Figure 3c). However, GSK2606414 reduced eIF2⍺ phosphorylation induced by all three PV antibodies, albeit only after 1 hour of PV exposure (Figure 3d). These findings demonstrate that miglustat and GSK2606414 prevent the PV IgG-induced ER stress response by targeting distinct pathways.

**Figure 3.**
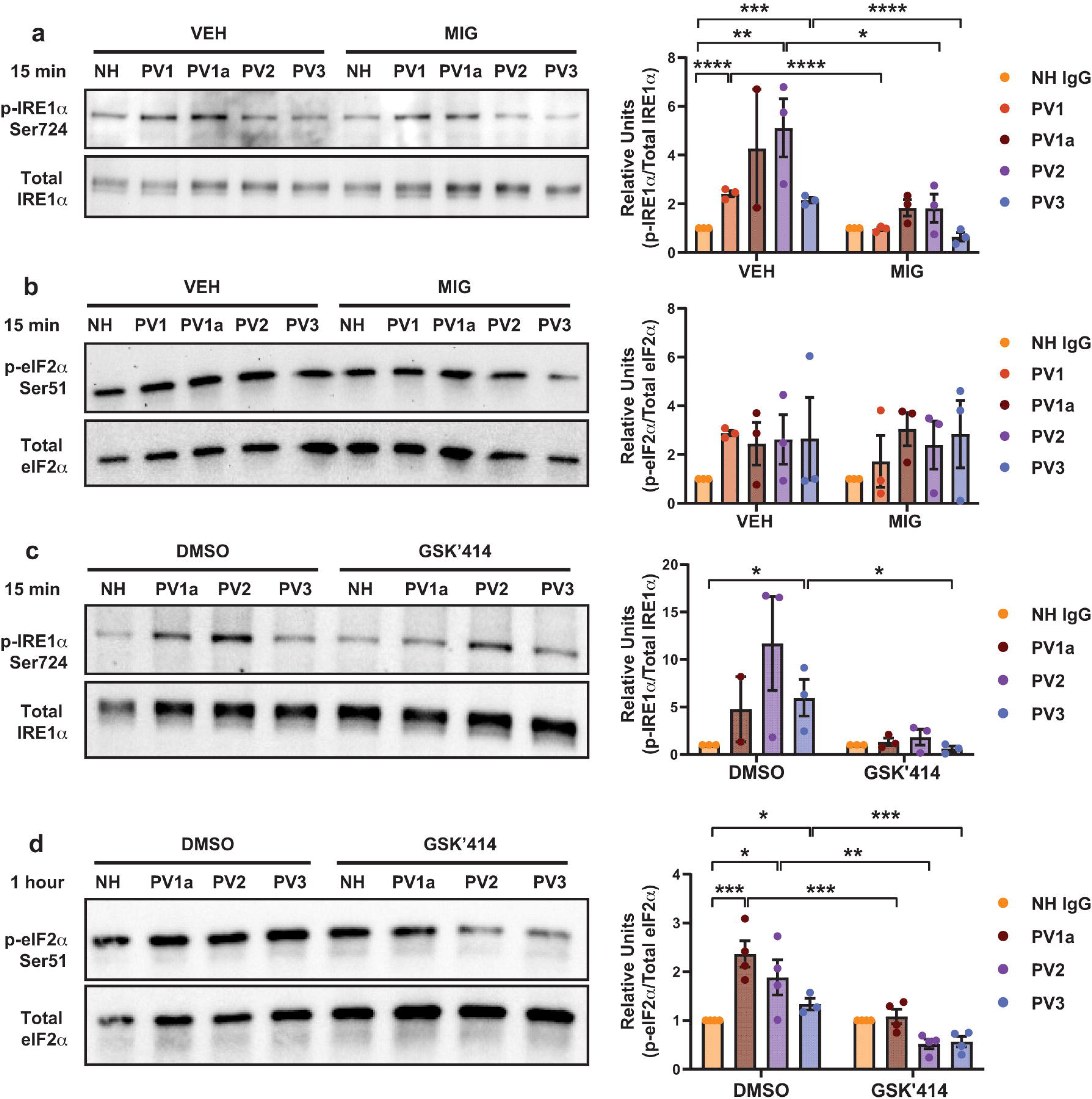
Miglustat and GSK2606414 prevent PV IgG-induced ER stress activation. NHEKs were pre-treated with vehicle control (VEH), miglustat (MIG), DMSO, or GSK2606414 (GSK’414) and then treated with IgG from NH or PV patients (PV1, PV1a, PV2, or PV3) for 15 minutes or 1 hour, and ER stress marker phosphorylation was monitored by western blot. **(a)** At 15 min, miglustat significantly prevented PV IgG-induced IRE1⍺ phosphorylation but **(b)** did not have a significant effect on eIF2⍺ phosphorylation. **(c)** GSK2606414 significantly prevented PV IgG-induced phosphorylation of IRE1⍺ phosphorylation at 15 minutes and **(d)** eIF2⍺ at 1 hour. Mean value ± SEM is represented. * *P* < 0.05, ** *P* < 0.01, *** *P* < 0.001, and **** *P* < 0.0001 using a two-way ANOVA.

To evaluate the effect of ER stress inhibition on PV IgG-induced desmosome disruption, we immunolabeled Dsg3 in NHEKs pre-treated with either miglustat or GSK2606414. Treatment with PV1a or PV2 dramatically disrupted Dsg3 border localization compared to cells treated with NH IgG. However, pre-treatment with either miglustat or GSK2606414 prevented PV IgG-induced disruption of Dsg3 border localization (Figure 4a). To assess the effect of ER stress inhibition on PV IgG-induced loss of cell-cell adhesion, we performed a dispase fragmentation assay (Hudson et al., 2004; Huen et al., 2002) in NHEKs pretreated with either miglustat or GSK2606414. PV IgG treatment caused extensive monolayer fragmentation compared to NH IgG. However, pre-treatment with either miglustat or GSK2606414 significantly reduced monolayer fragmentation in PV IgG-treated cultures (Figure 4b). Together, these results indicate that inhibition of ER stress protects against PV IgG-induced desmosome disruption and loss of cell-cell adhesion.

**Figure 4.**
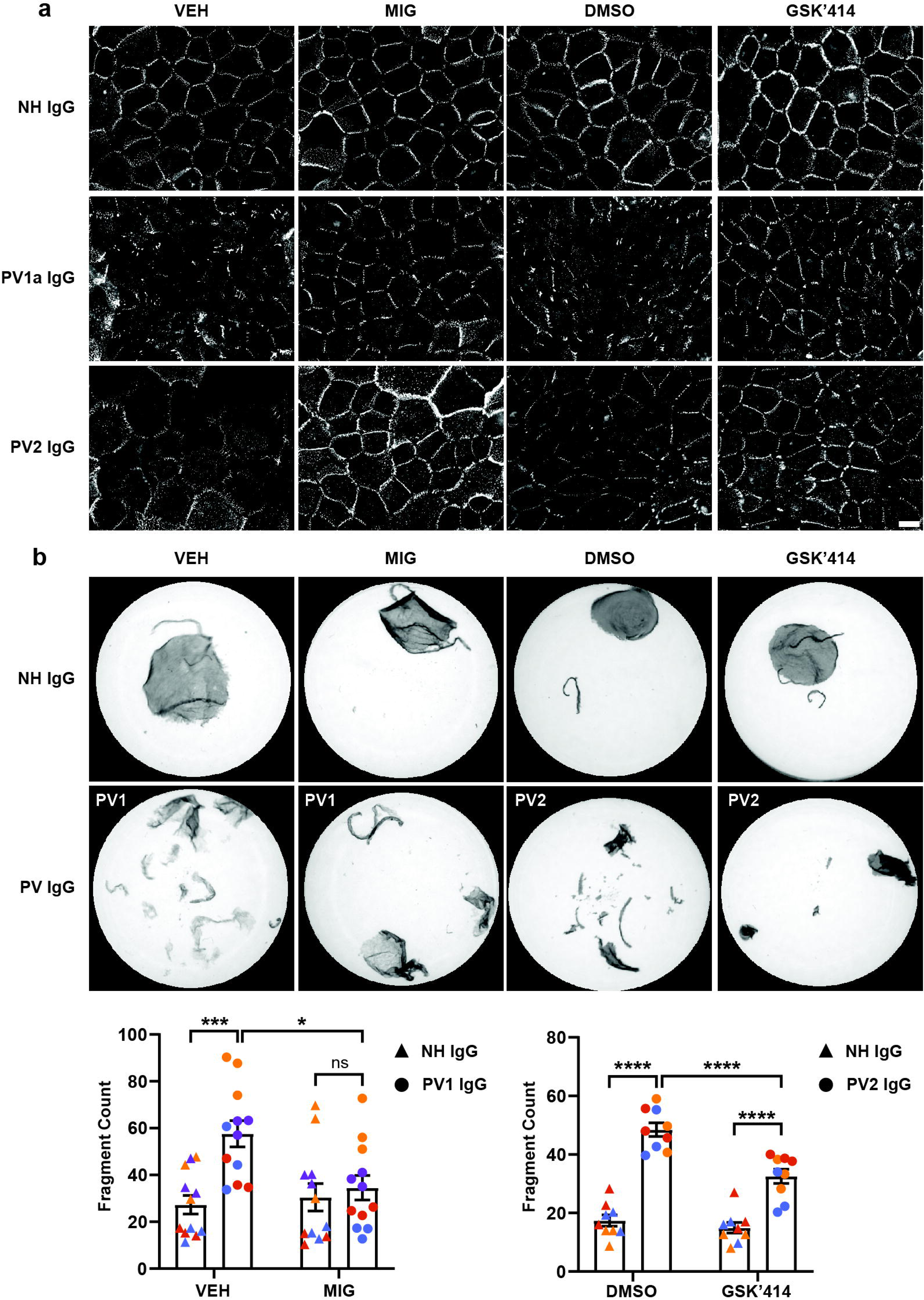
ER stress inhibitors prevent pathogenic PV IgG responses. NHEKs were pre-treated with vehicle control (VEH), miglustat (MIG), DMSO, or GSK2606414 (GSK’414) and then treated with IgG from NH or PV patients (PV1a or PV2). **(a)** Localization of Dsg3 was monitored by immunofluorescence microscopy. Dsg3 border localization was decreased by both PV IgG in the controls but not in miglustat- or GSK2606414-treated cells. Scale bar = 20 μm. **(b)** Loss of adhesion was assessed by dispase fragmentation assay. Fragmentation was significantly increased by PV IgG in the controls but not in miglustat- or GSK2606414-treated cells. Mean value ± SEM is represented. ns = not significant, * *P* < 0.05, *** *P* < 0.001, and **** *P* < 0.0001 using a two- way ANOVA.

## DISCUSSION

The results presented here demonstrate that the ER mediates a keratinocyte stress response to PV IgG. Live-cell confocal microscopy revealed that ER tubules associate with Dsg3 puncta as they internalize during desmosome disassembly in response to PV IgG. These findings are consistent with our previous work demonstrating that ER tubules form extensive contacts with both desmosomes and the keratin intermediate filament cytoskeleton (Figure 5) (Bharathan et al., 2023). ER stress signaling was observed in both cultured keratinocytes treated with PV IgG and in PV patient skin biopsies. Further, we demonstrate that pharmacological inhibition of ER stress alleviates the pathogenic effects of PV IgG on desmosome disruption and keratinocyte adhesion. Collectively, these findings suggest that the ER plays a role in the pathogenic mechanisms that underlie PV IgG-induced loss of adhesion.

**Figure 5.**
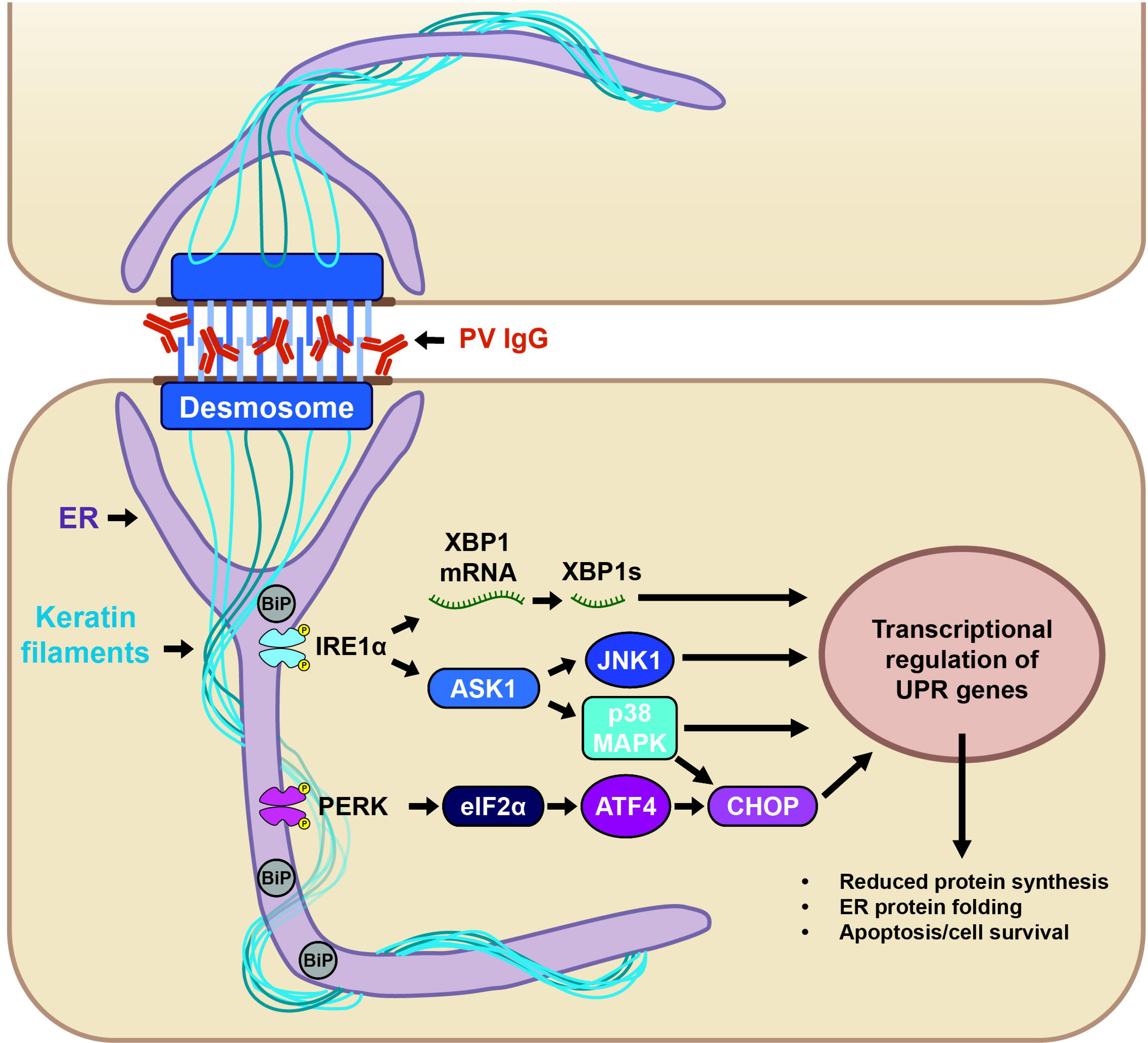
Proposed model of ER stress signaling in response to PV IgG. ER tubules (magenta) form close associations with the desmosome junction (dark blue) and keratin filaments (light blue) on either side of a cell-cell contact. ER-desmosome associations are maintained following PV IgG-mediated Dsg3 clustering and linear array formation. Exposure to PV IgG leads to activation of ER stress through the IRE1⍺ and PERK pathways, leading to transcriptional activation of the unfolded protein response and subsequent downstream signaling.

A functional link between ER and desmosome regulation first came from studies on Darier’s disease, a desmosomal skin disease that is caused by mutations in the ER calcium pump, sarcoendoplasmic reticulum Ca^2+^-ATPase isoform 2 (SERCA2) (Harmon et al., 2024 in press; Sakuntabhai et al., 1999). SERCA2 dysfunction in DD has been shown to cause dysregulation of desmoplakin dynamics, impaired adhesion strength and constitutive ER stress (Hobbs et al., 2011; Savignac et al., 2014; Sugiura, 2013). Further, inhibition of ER stress with chemical chaperones protects against keratin aggregation in epidermolysis bullosa simplex, a blistering skin disease (Chamcheu et al., 2011; Chamcheu et al., 2016). ER stress has also been implicated in the pathology of arrhythmogenic cardiomyopathy, a condition that can be caused by defective desmosomal adhesion (Pitsch et al., 2021). The activation of ER stress in these other diseases that display adhesion defects is similar to what we observe in PV, suggesting that mechanics and desmosomal adhesion are integrated with ER function and that skin disease models may be used to elucidate the functions of the ER-desmosome complex and its potential role in stress-sensing (Angulo-Urarte et al., 2020).

ER stress signaling intersects with various pathways that are known to be activated in PV, including p38 MAPK, PKC, and EGFR (Sharma et al., 2007; Spindler and Waschke, 2014; Takayanagi et al., 2015). p38 MAPK is a stress-activated protein kinase that is known to be involved in signaling both upstream and downstream of ER stress (Kim et al., 2008; Koeberle et al., 2015; Yang et al., 2007). p38 MAPK is phosphorylated as early as 15-30 min after PV IgG treatment in cultured keratinocytes (Berkowitz et al., 2005) and is also activated in PV patient skin (Berkowitz et al., 2008b). These findings are consistent with our results demonstrating that ER stress is rapidly activated by PV IgG (Figure 2), suggesting that p38 MAPK and ER stress pathways are functionally integrated. Further, p38 MAPK knockdown prevents PV IgG-induced loss of desmosomal Dsg3 in cultured keratinocytes (Mao et al., 2011) and p38 MAPK inhibition prevents blistering in mouse models of pemphigus (Berkowitz et al., 2008a; Berkowitz et al., 2006). Together with our observations, these findings suggest that ER stress may be a targetable pathomechanism in PV.

We used miglustat, an FDA-approved inhibitor of ceramide-specific glucosyltransferase that is thought to have molecular chaperone activities (Abian et al., 2011; Alfonso et al., 2005). In previous work, miglustat restored desmosome formation and improved adhesion strength in a keratinocyte model of Darier’s disease with constitutive ER stress (Savignac et al., 2014).

Consistent with the Savignac study, we show that miglustat did not affect eIF2⍺ phosphorylation. However, we find that miglustat does reduce IRE1⍺ phosphorylation by PV IgG (Figure 3). Interestingly, p38 MAPK, whose activation has been well-characterized in PV, is downstream of IRE1⍺, so it is possible that the IRE1⍺ pathway is the more prominent ER stress pathway that is activated in PV (Figure 5). Consistent with this hypothesis, miglustat had a significant protective effect against PV IgG-induced desmosome disruption and loss of adhesion (Figure 4). Overall, these findings suggest that ER stress pathways are activated in several types of desmosome disorders and may represent a therapeutically relevant pathway for treatment of desmosome diseases.

## METHODS

### Cell line generation, transfection, culture, and reagents

All cells were cultured at 37°C and 5% CO2. N/TERT keratinocytes were generated as previously described (Dickson et al., 2000). N/TERTs were cultured in Keratinocyte Serum-Free Media (K-SFM) (37010022, Gibco, Waltham, MA). For daily maintenance and subculturing of N/TERTs, the final calcium concentration was adjusted to 400 µM. The cells were switched to medium containing 550 µM calcium (high calcium media) 5-24 hours before experimentation.

N/TERTs used for live-cell imaging were transduced with a StayGold fluorescent protein as a soluble marker of the ER lumen (er-(n2)oxStayGold(c4)v2.0 or KDEL-StayGold) (Hirano et al., 2022) lentivirus by incubating cells with 8 µg/mL polybrene (TR-1003-G, EMD Millipore, Burlington, MA) in cell culture medium for 24 hours. Cells stably infected with the KDEL- StayGold lentivirus were selected using puromycin (4 μg/mL) (ABT-440, Boston BioProducts, Milford, MA). Bulk sorting of cell lines expressing the lentiviral constructs was performed by fluorescence-activated cell sorting to obtain populations with similar expression levels.

NHEKs were isolated from neonatal foreskin as previously described (Calkins et al., 2006; Schell et al., 2023). NHEKs (no later than passage 3) were cultured in KGM Gold Keratinocyte Growth Medium BulletKit (00192060, Lonza, Basel, Switzerland). For daily maintenance and subculturing of NHEKs, the final calcium concentration was adjusted to 50 µM to prevent desmosome formation. Cells were switched to high calcium media (550 µM) for 5-24 hours before experimentation to induce desmosome formation.

### Treatment of cells with PV IgG

Desmosome formation was induced by switching cells to high calcium media (550 µM) for 5-24 hours. Cells were treated with either normal healthy (NH) IgG or PV patient IgG (400 µg/mL) for indicated time periods at 37°C. PV sera were kind gifts from Dr. Masayuki Amagai (Keio University, Tokyo) and Dr. Aimee Payne (Columbia University, New York City, NY) (Saito et al., 2012; Stahley et al., 2016). All PV IgG used in this study cause loss of junctional Dsg3 as assessed by immunofluorescence, and loss of cell-cell adhesion as assessed by dispase fragmentation assay as shown previously for PV1 (Saito et al., 2012) or in this manuscript for PV1a, PV2, and PV3 (Figure 4, Supplemental Figure 1).

### Drug treatments

For miglustat (N-Butyldeoxynojirimycin) (B8299, Millipore-Sigma, Burlington, MA) experiments, cells were simultaneously switched to high calcium media (550 µM) and treated with either vehicle (sterile water) or 500 µM miglustat for indicated time periods. For GSK2606414 (HY-18072, MedChemExpress, Monmouth Junction, NJ) experiments, cells were switched to high calcium media (550 µM) for indicated time periods, then treated with either DMSO or 1 µM GSK2606414 for 1 hour.

### qRT-PCR analysis

RNA was isolated from previously collected and deidentified paraffin embedded PV patient biopsies using the Quick-RNA FFPE MiniPrep kit (ZR1008, Zymo, Irvine, CA). RNA concentration and purity were assessed using spectrophotometry. A total of 400 ng of RNA was reverse transcribed using the iScript cDNA Synthesis Kit (1708891, Bio-Rad, Hercules, CA).

For expression analysis of cells treated with normal human IgG or PV IgG, qRT-PCR was performed using TaqMan Fast Advanced Master Mix (4444557, Applied Biosystems, Waltham, MA) and Light Cycler 480 (Roche, Basel, Switzerland). TaqMan MGB probes labelled with fluorescent dyes were used. Reaction was performed according to the following protocol: 50°C for 2 min, 95°C for 20 sec, and (50 cycles of 95°C for 3 sec and 60°C for 30 sec).

Probes against target genes of interest were labelled with the FAM dye. *TBP* was labelled with the VIC dye and used as the reference gene within the same well as the target gene of interest. Normalized expression was calculated using Microsoft Excel (version 2023) by subtracting the cycle threshold (*Ct*) value of the internal control gene from the *Ct* value of the gene of interest, followed by averaging this value across all technical replicates. Fold change relative to healthy patient tissue was then calculated by the 2^-ΔΔCt^ method. More details about TaqMan probes are provided in Supplementary Table 1.

### Immunofluorescence

Cells were grown to 80% confluence on #1.5 glass coverslips coated with 0.1% gelatin solution (PCS-999-027, ATCC, Manassas, VA). For miglustat experiments, cells were simultaneously switched to high calcium media (550 µM) and treated with either vehicle (sterile water) or miglustat for 18 hours. For GSK2606414 experiments, cells were switched to high calcium media (550 µM) for 17 hours, and then treated with either DMSO or GSK2606414 for 1 hour. Drug-treated cells were then pre-labeled with P2C2, a non-pathogenic anti-Dsg3 monoclonal antibody conjugated to CF568 (P2C2-CF568) for 30 minutes at 4°C on a rocker, and then treated with NH or PV IgG in drug-containing high calcium media (550 µM) for 6 hours.

Cell borders were labeled with 2 µg/mL WGA-640R (29026, Biotium, Fremont, CA) in appropriate cell culture media at 37°C for 15 minutes. Cells were then washed three times with 1X PBS + 550 µM calcium for 5 minutes each, fixed with 100% methanol at -20°C for 2 minutes, followed by 3 more washes, and mounted with ProLong Glass Antifade Mountant (P36980, Invitrogen, Waltham, MA).

### Dispase fragmentation assay

Cells were grown to 100% confluence in 24-well plates. For miglustat experiments, cells were simultaneously switched to high calcium media (550 µM) and treated with either vehicle (sterile water) or miglustat for 24 hours. For GSK2606414 experiments, cells were switched to high calcium media (550 µM) for 5 hours, and then treated with either DMSO or GSK2606414 for 1 hour. Drug-treated cells were then treated with NH or PV IgG in drug-containing high calcium media (550 µM) for 18 hours. Monolayers were dissociated from the plate with dispase (354235, Corning, Corning, NY), using 40 µL dispase + 200 µL cell culture media per well in a 24-well plate. Monolayers were rinsed with 1X PBS + 550 µM calcium, transferred to 5 mL tubes, and subjected to mechanical stress on a tube rotator at 55 rpm for 1-3 min. Fragments transferred to 4-well plates were fixed and stained with 1% paraformaldehyde and methylene blue. Fragments imaged on a ChemiDoc MP Imaging System (Bio-Rad, Hercules, CA) were counted with Fiji.

## Data availability statement

Datasets related to this article are hosted on ScholarSphere (https://scholarsphere.psu.edu/resources/d8faef77-b085-44db-8610-5c5a7448070e), an open-source online data repository. Supplementary Videos S1-S3 can be accessed at: https://www.youtube.com/playlist?list=PLRVCHiGokrM4YVjJDkt_hH7ehM9V9iBPJ. All other data supporting the findings of this study are available from the corresponding author on reasonable request.

## ACKNOWLEDGMENTS

The authors thank Nate Sheaffer and Joseph Bednarczyk from Penn State College of Medicine’s Flow Cytometry Core for assistance with cell sorting. We are also grateful to the University of Pennsylvania Skin Biology and Diseases Resource-based Center (Penn SBDRC) as well as Amanda Nelson and Ryan Hobbs (Penn State College of Medicine) for instrument use, reagents, and advice. This work was supported by NIH grants R01AR081883 and R01AR048266 to APK, NIH P30-AR075043 to JEG, and NIH P30-AR-069589 to PI-Grice, Core Director JTS.

## AUTHOR CONTRIBUTIONS

Conceptualization: CLH, NKB, APK; Formal Analysis: CLH, NKB, WG; Funding Acquisition: APK; Investigation: CLH, NKB, YS; Resources: JEG, JTS, SMP, ASP; Software: NKB, WG; Visualization: CLH, NKB; Writing - Original Draft Preparation: CLH, NKB, APK; Writing - Review and Editing: CLH, NKB, APK

## SUPPLEMENTARY MATERIAL

**Supplementary Figure 1.**
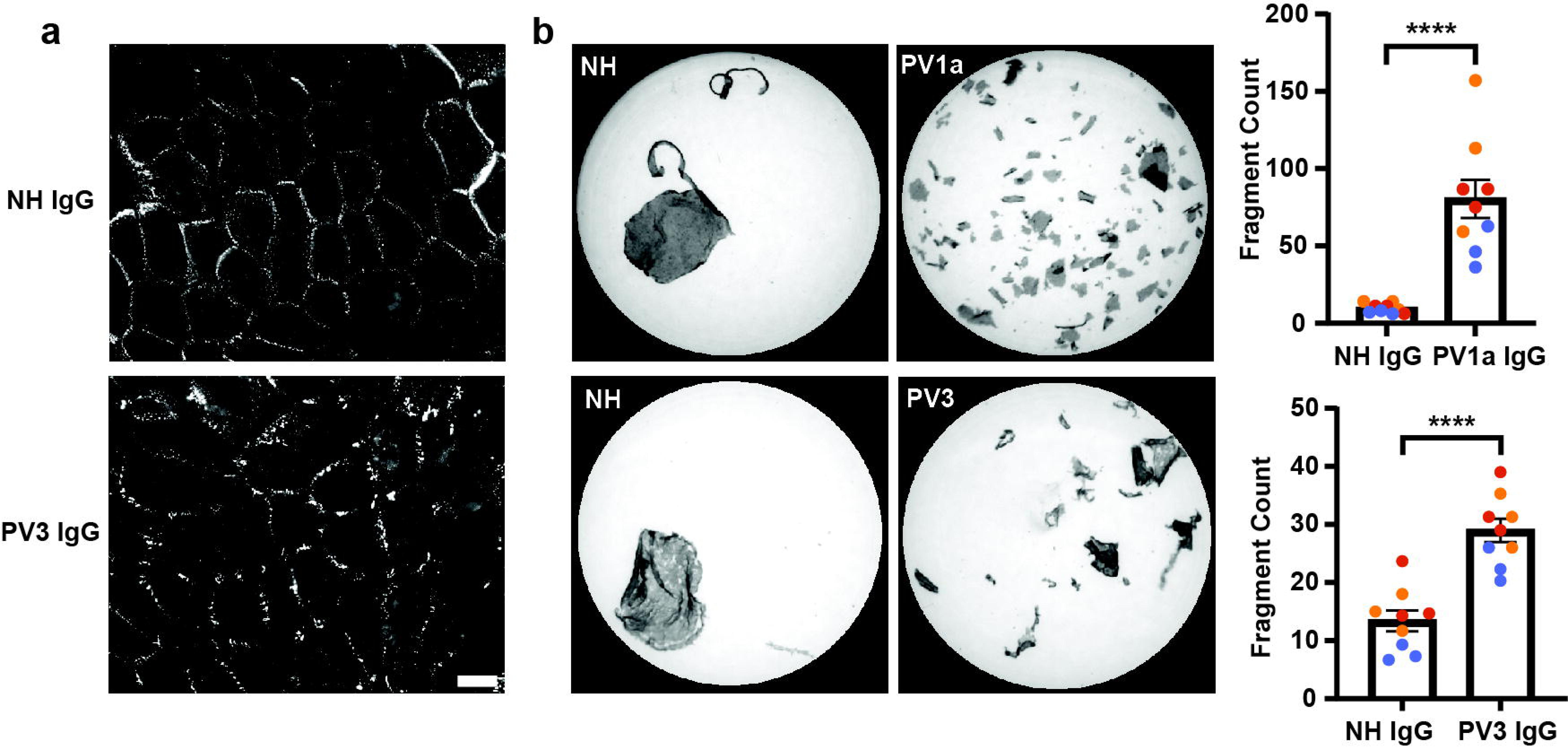
PV IgG cause loss of junctional Dsg3 and loss of cell-cell adhesion. NHEKs were switched to high calcium media (550 µM) for 6 hours and then treated with NH or PV IgG. **(a)** Localization of Dsg3 was monitored by immunofluorescence microscopy. Dsg3 border localization was decreased by PV3 IgG relative to NH IgG-treated cells. Scale bar = 20 μm. **(b)** Loss of adhesion was assessed by dispase fragmentation assay. Fragmentation was significantly increased by PV IgG relative to NH IgG-treated cells. Mean value ± SEM is represented. **** *P* < 0.0001 using an unpaired student’s t-test comparing PV IgG-treated to NH IgG-treated.

**Supplementary Video 1. ER tubules maintain stable associations with Dsg3 puncta in keratinocytes treated with NH IgG.** Live-cell spinning disk confocal microscopy of N/TERT keratinocytes treated with NH IgG shows that ER tubules (KDEL-StayGold, magenta) maintain associations with junctional Dsg3 puncta (anti-Dsg3 antibody, orange). Arrows point to ER tubules arranged in a mirror image-like pattern and stably anchored to Dsg3 puncta. Scale bar = 5 μm.

**Supplementary Video 2. ER tubules maintain stable associations with Dsg3 puncta in keratinocytes treated with PV IgG.** Live-cell spinning disk confocal microscopy of N/TERT keratinocytes treated with PV IgG shows that ER tubules (KDEL-StayGold, magenta) maintain associations with junctional Dsg3 puncta (anti-Dsg3 antibody, orange) during the first 30 minutes following PV IgG addition. Arrows point to ER tubules arranged in a mirror image-like pattern and stably anchored to Dsg3 puncta. Scale bar = 5 μm.

**Supplementary Video 3. ER tubules maintain stable associations with linear arrays in keratinocytes treated with PV IgG.** Live-cell spinning disk confocal microscopy of N/TERT keratinocytes treated with PV IgG shows that ER tubules (KDEL-StayGold, magenta) maintain associations with linear arrays of Dsg3 (anti-Dsg3 antibody, orange) seen forming perpendicular to the cell-cell contact. Arrows point to ER tubules maintaining stable associations with linear arrays over time. Scale bar = 5 μm.

**Supplementary Table 1.**
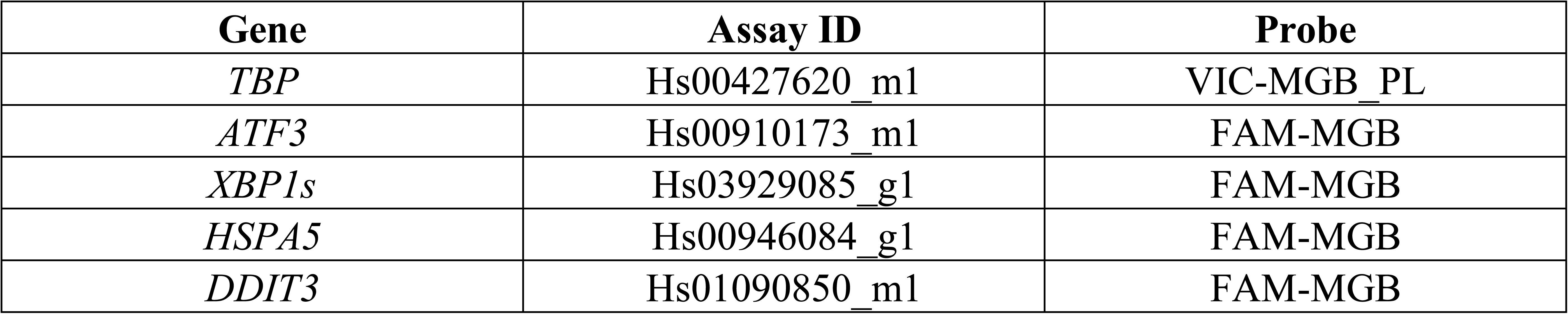
TaqMan probes used in this study.

**Supplementary Table 2.** Spinning disk confocal acquisition settings.

| Figure | Number of slices | Z-step size [μm] | Exposure time [ms] | Interval [s] | Duration [min] |
| --- | --- | --- | --- | --- | --- |
| Figure 1a | 7 | 0.45 | 50 | 10 | 20 |
| Figure 1b | 7 | 0.45 | 50 | 10 | 5 |
| Figure 1c | 7 | 0.45 | 50 | 10 | 10 |

## SUPPLEMENTARY METHODS

### Lentivirus generation

pcDNA3/er-(n2)oxStayGold(c4)v2.0, a KDEL-StayGold sequence codon-optimized for mammalian expression, was a gift from Atsushi Miyawaki (Addgene plasmid #186296) (Hirano et al., 2022). A lentiviral plasmid containing this KDEL-StayGold sequence was synthesized by VectorBuilder (VectorID: VB220722-1159spe). Whole plasmid sequencing of the KDEL- StayGold plasmid was performed by Plasmidsaurus using Oxford Nanopore Technology with custom analysis and annotation. Lentiviruses were made by co-transfection of lentiviral plasmids, pMD2.G (encoding VSV-G), and psPAX2 (encoding Gag and Pol) into human embryonic kidney-293FT cells, followed by collection of culture supernatants 24-72 hours after transfection. Lentivirus was concentrated with the Lenti-X Concentrator kit (631231, Takara Bio, Kasatsu, Shiga, Japan) following the manufacturer’s protocol. The pMD2.G and psPAX2 (gifts from Didier Trono, plasmid #12259 and #12260, respectively) plasmids were obtained from Addgene. Plasmid sequences are available at https://doi.org/10.5281/zenodo.13138745.

### Western blotting

Cells were grown to 90% confluence in 12-well plates. For miglustat experiments, cells were simultaneously switched to high calcium media (550 µM) and treated with either vehicle (sterile water) or miglustat for 6 hours. For GSK2606414 experiments, cells were switched to high calcium media (550 µM) for 5 hours, and then treated with either DMSO or GSK2606414 for 1 hour. Cells were then treated with NH or PV IgG in drug-containing high calcium media (550 µM) for various time periods and then incubated with 1X Laemmli buffer (1610737, Bio-Rad, Hercules, CA) with β-mercaptoethanol (M2650, Sigma-Aldrich, St. Louis, MO) for 5 minutes. Lysates were then collected and incubated at 95-100°C for 10 minutes. Proteins were separated by SDS-PAGE along with a protein ladder (26634, Thermo Fisher Scientific, Waltham, MA) using Tris/glycine/SDS running buffer (161-0732, Bio-Rad, Hercules, CA). When probing for IRE1⍺, proteins were separated on 7.5% Mini-PROTEAN TGX Gels (4561026, Bio-Rad, Hercules, CA) and then transferred using 2X transfer buffer (161-0734, Bio-Rad, Hercules, CA) containing 20% methanol. Transfer to a nitrocellulose membrane (88018, Thermo Fisher Scientific. Waltham, MA) was performed overnight at 22 V. When probing for eIF2⍺, proteins were separated on 10% Mini-PROTEAN TGX Gels (4561036, Bio-Rad, Hercules, CA) and transferred using 1X Trans-Blot Turbo Transfer Buffer (10026938, Bio-Rad, Hercules, CA). Transfer to a nitrocellulose membrane was performed using the Trans-Blot Turbo Transfer System for 7 minutes at 25 V. Membranes were blocked in 2.5% milk (in 1X TBS-Tween) for 1 hour. Membranes were incubated with primary antibodies (in 5% BSA in 1X TBS-Tween) at 4°C overnight. All blots were probed for ⍺-tubulin to ensure equivalent protein loading. Membranes were washed 3 times with 1X TBS-Tween for 5 minutes each, followed by incubation in secondary antibody for 1 hour, followed by 3 more washes (5 minutes each).

Membranes were always probed for phosphorylated antibodies first and then stripped with OneMinute Western Blot Stripping Buffer (GM6001, GM Biosciences, Frederick, MD), followed by a 30-minute block. The above wash and incubation steps were then repeated for the next primary and secondary antibodies. Western blots were developed with chemiluminescence horseradish peroxidase substrate (RPN2232, Cytiva, Marlborough, MA). The chemiluminescent blots were imaged with a ChemiDoc MP Imaging System (Bio-Rad, Hercules, CA).

Densitometric analysis was performed using Fiji/ImageJ (v2.14.0/1.54f) (Schindelin et al., 2012).

### Antibodies

The following antibodies were used for western blot: 1:1000 rabbit anti-phospho-eIF2⍺ (pSer51) (CS#3398, Cell Signaling, Danvers, MA), 1:1000 rabbit anti-eIF2⍺ (CS#9722, Cell Signaling, Danvers, MA), 1:1000 rabbit anti-phospho-IRE1⍺ (pSer-724) (NB100-2323, Novus Biologicals, Centennial, CO), 1:500 rabbit anti-IRE1⍺ (14C10) (CS#3294, Cell Signaling, Danvers, MA), 1:1000 mouse anti-⍺-tubulin (12G10) (Iowa Hybridoma Bank, Iowa City, Iowa). Secondary antibodies used were 1:3000 goat anti-rabbit IgG (H + L) horseradish peroxidase conjugate (170-6515, Bio-Rad, Hercules, CA), and 1:3000 goat anti-mouse IgG (H + L) horseradish peroxidase conjugate (170-6516, Bio-Rad, Hercules, CA). For fixed and live-cell immunofluorescence, we conjugated the fluorescent tag CF568 to a patient-cloned non-pathogenic monoclonal antibody against Dsg3 (P2C2-568) using Mix-n-Stain CF Dye Antibody Labeling Kit (92447, Biotium, Fremont, CA).

### Spinning disk confocal microscope

Fluorescence imaging was performed using a Nikon Ti2-E equipped with a Yokogawa CSU-X1 spinning disk unit, LUNF XL laser unit, Nikon Perfect Focus System, Z piezo stage, motorized XY stage, two sCMOS cameras (ORCA-Fusion BT, Hamamatsu Corp., Bridgewater, NJ), and two fast filter wheels with most elements controlled through hardware-triggering through the Nikon’s National Instruments Breakout Box. The acquisition software was NIS-Elements (v5.42.03). The polychroic mirror within the Yokogawa CSU-X1 unit is a Semrock Di01-T405/488/568/647. Single emission filters (Chroma ET525/36m, Chroma ET605/52m, and Chroma ET705/62m) were used with the 488nm, 561nm, and 640nm lasers.

### Widefield microscope

Fixed immunofluorescence images were acquired using a Zeiss AxioObserver microscope with the ZEN acquisition software (version 2.3). We used a Zeiss Plan-APOCHROMAT 63x/1.4 NA oil DIC immersion objective and an X-Cite 120 LED Boost with the following filters: Zeiss filter set 43 HE (excitation: BP 550/25 (HE), beamsplitter: FT 570 (HE), emission: BP 605/70 (HE)) for CF-568 and Zeiss filter set 50 (excitation: BP 640/30, beamsplitter: FT 660, emission: BP 690/50) for CF-640R. Images were captured with an Axiocam 506 monochrome camera.

### Live-cell imaging

For live-cell imaging, cells were seeded on 8-well chambered #1.5H cover glass (#C8- 1.5H-N, Cellvis, Mountain View, CA). We used a Nikon 100x/1.49 NA Apo TIRF oil immersion objective with its correction collar optimized for our imaging at 37°C. Images were taken in 12- bit with high gain (“12-bit Sensitive”) and with “Standard” readout mode. Rough alignment for simultaneous dual-channel imaging was accomplished by using the spinning disk pinholes in the OFF position to align the Cairn TwinCam unit (holding a Semrock Di02-R561 beamsplitter) until pixel-perfect overlap was achieved in the center of FOV for the two cameras. Temperature and CO2 control for maintaining physiological conditions were provided by a Tokai Hit stage top incubation system (Model STXF-WELSX-SET). Detailed acquisition settings for figures/experiments are listed in Supplementary Table 2.

To image the interaction between the ER and desmosomes in living cells, N/TERT keratinocytes stably expressing KDEL-StayGold were incubated in K-SFM (containing 550 µM calcium) for 18-24 hours to induce desmosome formation. The next day, Dsg3 was pre-labeled by incubating cells with P2C2-568 in cold (4°C) K-SFM (containing 550 µM calcium) for 30 minutes at 4°C on a low-speed orbital shaker. Cells were then washed twice with 1X PBS+ (containing 550 µM calcium), followed by addition of warm K-SFM (containing 550 µM calcium) with 400 µg/ml NH or PV IgG. The 8-well dish was then immediately placed on the confocal microscope for live-cell imaging.

### Fluorescence microscopy image processing

Images were split by channels and timepoints before denoising with Noise2Void (Krull et al., 2019). After re-combining to a hyperstack, all datasets were corrected for lateral chromatic aberration using NanoJ’s channel registration (Laine et al., 2019). If needed, images were then drift corrected using NanoJ’s drift correction. Fiji/ ImageJ macros are provided at https://doi.org/10.5281/zenodo.13138745.

### Noise2Void

Noise2Void v0.2.1 was used with TensorFlow-DirectML on a Windows 10 workstation with an NVIDIA RTX 3090 GPU (Graphics Processing Unit).

### Statistics

Parametric methods including Student’s unpaired two-tailed *t*-test (when two groups were compared) or two-way ANOVA with Tukey’s post hoc test (when >2 groups were compared with more than 1 independent variable), were used (GraphPad Prism 10.2.3, GraphPad Software, San Diego, CA). Results were considered significant when *P* < 0.05.

### Study approval

Experiments using discarded and/or de-identified human skin and pemphigus patient sera were reviewed and determined to be “not human subjects research” by the institutional review board at the Pennsylvania State University Office for Research Protections (PSU IRB STUDY00021792).

